# Root caries on a *Paranthropus robustus* third molar from Drimolen

**DOI:** 10.1101/573964

**Authors:** Ian Towle, Alessandro Riga, Joel D. Irish, Irene Dori, Colin Menter, Jacopo Moggi-Cecchi

**Affiliations:** Research Centre in Evolutionary Anthropology and Palaeoecology, School of Natural Sciences and Psychology, Liverpool John Moores University, Liverpool, United Kingdom, L3 3AF; Department of Biology, University of Florence, Italy; Evolutionary Studies Institute and Centre for Excellence in PaleoSciences, University of the Witwatersrand, Private Bag 3, WITS 2050, South Africa; Centre for Anthropological Research, University of Johannesburg, Johannesburg, South Africa

**Keywords:** caries, dental pathology, hominins, South Africa

## Abstract

**Objectives:** Dental caries is often perceived as a modern human disease. However, their presence is documented in many early human groups, various non-human primates and, increasingly, our hominin ancestors and relatives. In this study we describe an antemortem lesion on the root of a *Paranthropus robustus* third molar from Drimolen, South Africa, which likely represents another example of caries in fossil hominins.

**Materials and Methods:** The molar, DNH 40, is dated to 2.0–1.5 Ma and displays a lesion on the mesial root surface, extending from the cementoenamel junction 3 mm down toward the apex. The position and severity of the lesion was macroscopically recorded and micro-CT scanned to determine the extent of dentine involvement.

**Results:** A differential diagnosis indicates root caries, as the lesion is indistinguishable from clinical examples. Although necrotic in appearance, external tertiary dentine is evident on a micro CT scan. Gingival recession and/or continuous eruption of the tooth as a result of extensive occlusal wear would have occurred to facilitate caries formation. Therefore, the lesion is likely linked to relative old age of this individual.

**Discussion:** This new example increases the total number of carious lesions described in *P. robustus* teeth to 12, on occlusal, interproximal and, now, root surfaces. Beyond the consumption of caries-causing food(s), caries formation would have also required the presence of requisite intra-oral cariogenic bacteria in this individual and the species. Of interest, the presence of tertiary dentine on the outward surface suggests the DNH 40 lesion may have been arrested, i.e., no longer active, perhaps relating to a change in diet or oral microbiome just prior to the individual’s death.

## Introduction

Carious lesions may form when intraoral bacteria, notably *Streptococcus mutans* and *Lactobacillus* sp., excrete acidic waste as they metabolise ingested sugars and starches (Nishikawara et al., 2007). As the local pH lowers these bacteria can proliferate, leading to demineralization of dental tissues (Gussy et al., 2006). Foods containing refined carbohydrates and sugars are particularly cariogenic (Rohnbogner & Lewis, 2016); some fruits and nuts, as well as honey, may be as well (Novak, 2015; Humphrey et al., 2014). Conversely, fibrous/tough dietary items, meat and certain plants, among others, inhibit lesion formation (Moynihan, 2000; Rohnbogner & Lewis, 2016; Novak, 2015). As such, the absence, presence and characteristics of caries can give insight into the diet and food processing behaviours of individuals and groups (Takahashi & Nyvad, 2016).

Caries is often perceived as a disease of recent humans, that is rare or absent in fossil hominins (e.g., Tillier et al., 1995). However, carious lesions are well known in the archaeological record, particularly among agricultural groups, and there is a growing number recorded in various fossil hominin and nonhuman species (e.g., Lacy, 2014; Towle, 2017; Arnaud et al., 2016; Trinkaus et al., 2000; Margvelashvili et al., 2016). In this study we describe a root lesion in a tooth assigned to *Paranthropus robustus* from Drimolen (Gauteng, South Africa) (Moggi-Cecchi et al., 2010). A differential diagnosis is then presented to characterize this lesion and, further, assess the diet and state of oral health of the individual.

## Materials and methods

The tooth, DNH 40, is a left maxillary third molar (Moggi-Cecchi et al., 2010) dated to 2.0–1.5 Ma (Keyser et al., 2000; Pickering et al., 2018). A full morphological description is provided in Moggi-Cecchi et al. (2010). For this study, the position, extent and severity of the lesion was macroscopically observed and recorded. It was then micro-CT scanned to determine the extent of dentine involvement, i.e., lower tissue density from carious activity vs. higher density that potentially identifies tertiary dentine formation (Carvalho and Lussi, 2017). Other potential lesion-forming processes are also discussed and considered.

## Results and discussion

The 7 x 3 mm lesion extends from the cementoenamel junction (CEJ) on the tooth’s mesial root surface almost a quarter of the way down toward the apex (Figure 1A/B). Its shape, depth, and position is analogous to clinical and archaeological examples of root caries. Non-carious cervical lesions from abrasion, erosion or abfraction can appear superficially similar to root caries (Grippo et al., 2018; Towle et al., 2018). However, the lesion is neither consistent with an aetiology of abfraction nor erosion, in that it lacks both a smooth surface and ‘wedge-shaped’ appearance (Litonjua et al., 2003). Abrasion is also unlikely, given the lesion’s depth, shape and position in the overall dentition (Grippo et al., 2018).

**Figure 1.**
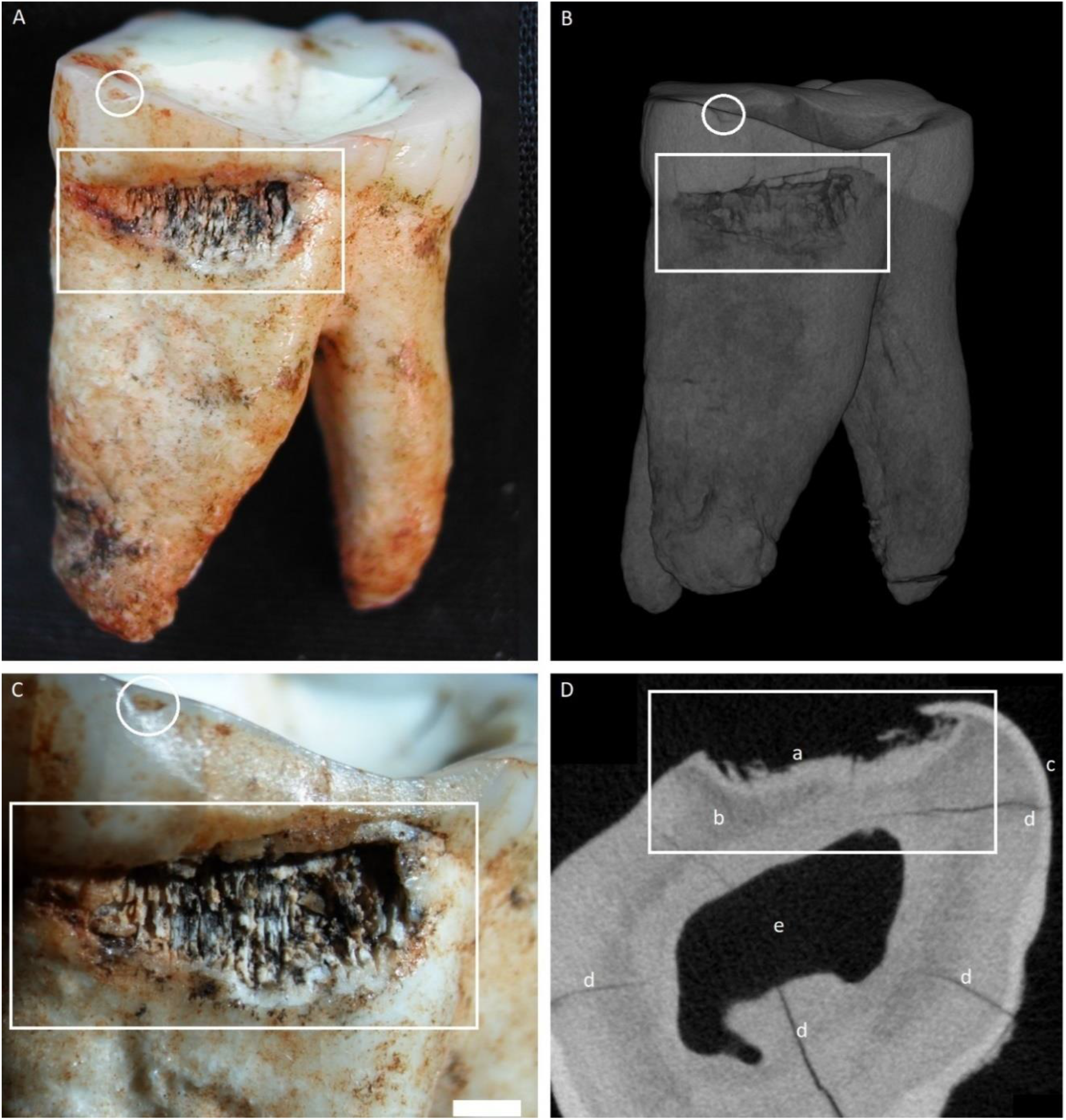
Specimen DNH 40, an upper left *Paranthropus robustus* third molar. White squares indicate the carious lesion, and white circles highlight an antemortem enamel fracture. A) Overview of the whole tooth, showing mesial and occlusal surfaces; B) Micro-CT rendering of the specimen; C) Close-up of the lesion, white bar is 1mm.; D) CT slice of the specimen, a: Tertiary dentine (light band of higher density), b: primary dentine, c: cementum, d: post-mortem cracks, e: pulp chamber.

It can be seen that the inferior enamel margin was subject to some carious activity (Figure 1C), but the dentine was primarily affected. That said, the pulp chamber was not perforated, while signs of tertiary dentine formation are evident on the micro-CT scan (Figure 1D). The carious lesion had formed in the interproximal area directly above the contact wear facet with the second molar (Figure 1A). An area of lower density primary dentine is evident directly behind the tertiary dentine band (Figure 1D, dark area around ‘b’), which likely relates to initial tissue demineralisation. The presence of tertiary dentine suggests the lesion was arrested, i.e., in the process of ‘healing’ at the time the individual died (Schüpbach et al., 1992; Carvalho and Lussi, 2017).

The lesion described here adds to the total number, as well as types of caries reported so far in fossil hominins—in this case 12 in *P. robustus* alone (Clement, 1956; Grine et al., 1990; Robinson, 1952; Towle, 2017). Because caries formation requires fermentable carbohydrates and cariogenic intra-oral bacteria (Clarkson et al., 1987), it is evident that both were present for at least the past 2.0-1.5 million years (Arnaud et al., 2016; Lacy, 2014; Trinkaus et al., 2000; Towle, 2017; Margvelashvili et al., 2016). Hominin species with caries frequencies below 1% or above 5% of teeth usually have a specific behavioral explanation for the rate (Towle, 2017). This is the case for many agricultural and hunter-gatherer diets, with specific cariogenic foods or tough non-cariogenic vegetation/meat (e.g., Novak, 2015; Vodanović et al., 2005; Slaus et al., 2011; Srejic, 2001; Varrela, 1991; Humphrey et al., 2014; Kelley et al., 1991; Lacy, 2014; Larsen et al., 1991). There is growing evidence that this relatively high caries rate is particularly characteristic of the *Homo* genus (e.g., Arnaud et al., 2016; Lacy, 2014; Lanfranco & Eggers, 2012; Lacy et al., 2012; Liu et al., 2015; Margvelashvili et al., 2016). Given that 12 caries have been described in *P. robustus,* it is likely that their diet was more similar to earlier *Homo* than once thought, at least to the extent that both consumed cariogenic foods. In contrast, caries has not yet been reported in *Australopithecus africanus,* though the latter sample size is similar to that of *P. robustus.* Consequently, this study adds further evidence to suggest these latter two species had significantly different diets.

With specific reference to root caries, formation is contingent upon access by cariogenic bacteria to the affected tissue through either 1) alveolar resorption, often from periodontitis in modern examples and/or 2) continuous eruption (super-eruption) of teeth in compensation of heavy occlusal wear (Hillson, 2008). In the present case, the latter factor is likely due to the wear on the occlusal surface (Figure 1A). Beyond the obvious, root exposure is further suggested by excess cementum deposition (aka. hypercementosis) near the root apices (Figure 1B), which can serve to help anchor the tooth in the absence of a complete alveolus.

The DNH 40 specimen displays a typical root caries pattern, i.e., spread out on the interproximal root surface and does not deeply penetrate the dentine (Carvalho and Lussi, 2017; Damé-Teixeira et al., 2017; Grippo et al., 2018). Grippo et al. (2018) suggest that stress associated with mastication could be intensified at interproximal sites, which may contribute to caries formation there. The molar not only exhibits heavy occlusal wear, but deep cupping on the mesial occlusal surface and an antemortem chip below the caries (Figure 1A/C). Therefore, high occlusal loads or regular consumption of tough foods may have contributed to the formation—in terms of interproximal stress and continuous eruption, which facilitated root exposure. Relative old age of the individual was likely a contributing factor to the carious lesion, since DNH 40 is a well-worn third molar and both gingivitis-related recession and continuous eruption are associated with aging (Hayes et al., 2017; Hillson, 2008). Some studies in humans suggest that root caries occur more often in females because of earlier alveolar resorption (Temple, 2016; Watson et al., 2010); however, such a link cannot be made for DNH 40 as the individual’s sex is unknown.

Root caries is rare in fossil hominins, likely because they require the root exposure in the oral cavity, and may be formed by a different mix of bacteria or in different acidic conditions (Shen et al., 2005). This study adds additional evidence to suggest that caries of all types affected other hominin species, in some cases with frequencies similar to those of recent humans. A likely explanation for the lesion in the present study was consumption of cariogenic fruits or vegetal matter, such as certain tubers; given the increasing number of severe lesions noted in *P. robustus,* consumption of honey may also be plausible. The root caries progression seemingly was arrested at the time of death, which may have resulted from a change in diet, a change in the local microbiome, loss of the adjacent second molar and/or seasonal changes (Miles & Grigson, 2003). In any event, the caries described here, whatever its etiology and state of ‘healing,’ provides further evidence that, among other hominin species, *P. robustus* suffered from dental problems not unlike those of modern humans.

## Acknowledgments

For CT scanning assistance we thank Jean-Jacques Hublin and Matthew Skinner, Department of Human Evolution, Max Planck Institute for Evolutionary Anthropology. Work at the Drimolen site by the Italian Archaeological Mission was supported by a series of grants (2006 – 2016) by the Italian Ministry of Foreign Affairs to JM-C. CM thanks the National Research Foundation (African Origins Platform) for grants that supported the excavation and research at Drimolen.

